# Ophiobolin A Covalently Targets Complex IV Leading to Mitochondrial Metabolic Collapse in Cancer Cells

**DOI:** 10.1101/2023.03.09.531918

**Authors:** Flor A. Gowans, Danny Q. Thach, Yangzhi Wang, Belen E. Altamirano Poblano, Dustin Dovala, John A. Tallarico, Jeffrey M. McKenna, Markus Schirle, Thomas J. Maimone, Daniel K. Nomura

**Affiliations:** Department of Nutritional Sciences and Toxicology, University of California, Berkeley, Berkeley, CA 94720 USA; Department of Chemistry, University of California, Berkeley, Berkeley, CA 94720 USA; Novartis-Berkeley Translational Chemical Biology Institute, Berkeley, CA 94720 USA; Innovative Genomics Institute, Berkeley, CA 94704 USA; Novartis Institutes for BioMedical Research, Emeryville, CA 94608 USA; Novartis Institutes for BioMedical Research, Cambridge, MA 02139 USA; Novartis Institutes for BioMedical Research, Basel, Switzerland; Department of Molecular and Cell Biology, University of California, Berkeley, Berkeley, CA 94720 USA

**Keywords:** chemoproteomics, activity-based protein profiling, ophiobolin, cysteine, lysine, mitochondrial metabolism, natural products

## Abstract

**Summary:** Ophiobolin A (OPA) is a sesterterpenoid fungal natural product with broad anti-cancer activity. While OPA possesses multiple electrophilic moieties that can covalently react with nucleophilic amino acids on proteins, the proteome-wide targets and mechanism of OPA remain poorly understood in many contexts. In this study, we used covalent chemoproteomic platforms to map the proteome-wide reactivity of OPA in a highly sensitive lung cancer cell line. Among several proteins that OPA engaged, we focused on two targets—cysteine C53 of HIG2DA and lysine K72 of COX5A—that are part of complex IV of the electron transport chain and contributed significantly to the anti-proliferative activity. OPA activated mitochondrial respiration in a HIG2DA and COX5A-dependent manner, led to an initial spike in mitochondrial ATP, but then compromised mitochondrial membrane potential leading to ATP depletion. We have used chemoproteomic strategies to discover a unique anti-cancer mechanism of OPA through activation of complex IV leading to compromised mitochondrial energetics and rapid cell death.

## Introduction

Natural products isolated from microbes, plants, and other living organisms have been a major source of therapeutics; they, and their derivatives, constitute 50-70% of the drugs used for the treatment of bacterial and fungal infections, inflammation, diabetes, and cancer ^1–4^. Among bioactive natural products there exist a subset of molecules that possess reactive electrophilic centers that can covalently react with nucleophilic amino acid residues on proteins, thus imparting therapeutic activity ^5^. Examples of such molecules include: *i)* lipstatin and penicillin which feature reactive *β*-lactone or -lactam warheads; *ii)* anti-cancer agents epoxomicin and fumagillin that harbor reactive epoxides; and *iii)* the anti-cancer agent Wortmannin that covalently modifies PI3 kinase ^3,6^. Beyond these examples, there are many other electrophilic natural products that show anti-cancer activity but their direct targets and mechanisms are poorly understood.

Ophiobolin A (OPA) is a phytotoxic fungal-derived sesterterpenoid natural product that has been shown to possess anti-proliferative activity across dozens of cancer cell types, often with sub-micromolar potency **(Figure 1a)** ^7^. OPA was originally isolated from the fungus *Ophiobolus miyabeanus* in 1958 by Ishibashi and Nakamura and its structure and stereochemistry were reported in 1965 ^7^. While OPA was initially characterized as a calmodulin inhibitor in plants ^8,9^, more recent studies have demonstrated that OPA impairs breast cancer, leukemia, glioma, and glioblastoma viability among other tumor cell lines ^1^. These findings have made ophiobolin sesterterpenes attractive targets for total synthesis and chemical probe development ^10–14^. The cytotoxic effects of various ophiobolin members have been attributed to the perturbation of numerous biological molecules, cell signaling nodes, and cell death pathways ^15–25^, and are suggestive of context-specific mechanisms of cell killing ^24^. Given OPA’s known ability to engage lysine residues of calmodulin in Schiff base formation ^8^, covalent chemistry – involving both lysine and cysteine – has also been implicated in OPA bioactivity ^25–28^.

**Figure 1.**
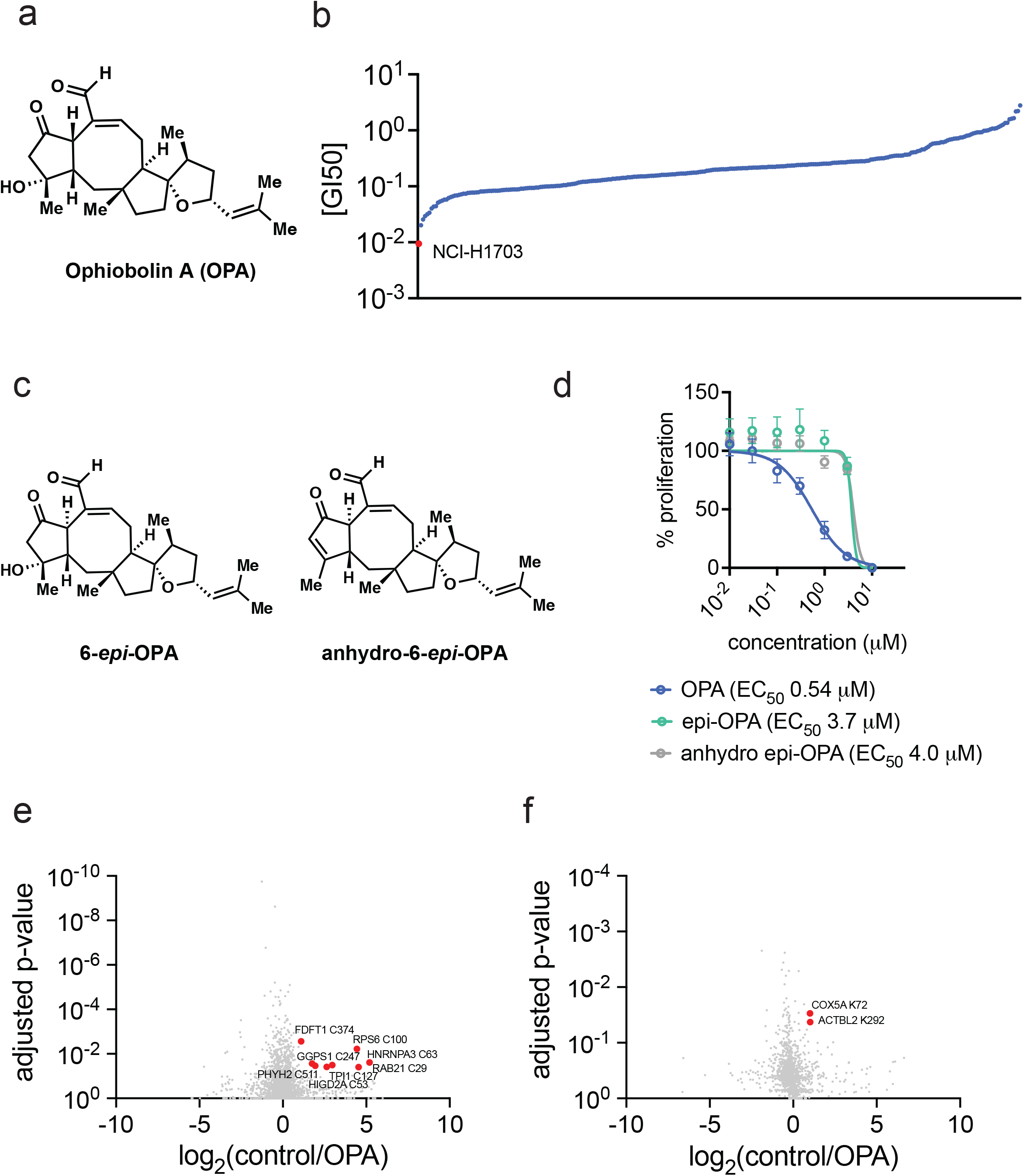
Characterizing OPA anti-cancer activity and proteome-wide cysteine and lysine reactivity. **(a)** OPA’s structure. OPA possesses a reactive aldehyde and a Michael acceptor that can potentially react with nucleophilic amino acids such as cysteine or lysine within protein targets. **(b)** 50 % growth inhibitory (GI_50_) values of OPA across 283 human cancer cell lines showing that NCI-H1703 was the most sensitive cell line to OPA with a GI_50_ of 0.17 µM. **(c)** Structures of OPA analogs 6-*epi*-OPA and anhydro-6-*epi*-OPA. **(d)** Cell proliferation in NCI-H1703 lung cancer cells treated with DMSO vehicle, OPA, 6-*epi*-OPA and anhydro-6-*epi*-OPA for 24 h, assessed by Hoechst stain. **(e, f)** Proteome-wide cysteine **(e)** and lysine **(f)** reactivity of OPA. NCI-H1703 cells were treated with DMSO vehicle or OPA (3 µM) for 4h. Subsequent cell lysates were treated with the cysteine-reactive alkyne-functionalized iodoacetamide probe (IA-alkyne) **(e)** or the lysine-reactive alkyne-functionalized NHS-ester probe (NHS-ester-alkyne) **(f)** and subjected to the isoTOP-ABPP protocol. Shown are control versus OPA probe-modified peptide ratios and adjusted p-values. Shown in red are the modified residue and targets that showed >2-fold control/OPA ratio with p<0.05. Data in **(b)** can be found in **Table S1**. Data shown in **(d)** are in average ± sem, n=5 biologically independent samples/group. Data shown in **(e, f)** are average probe-modified peptide ratios with n=3 biological independent samples/groups. Related to **Figure S1, Table S1, Table S2**, and **Table S3**.

Various studies have observed that OPA causes morphological and functional changes to mitochondria although the detailed mechanisms underlying these perturbations are unclear ^19,20,29^. One potential mechanism that has been invoked is the covalent inhibition of human calmodulin by OPA ^18,30^. Another more thorough mechanistic study of OPA using a functional genomic screen revealed that reducing de novo synthesis of the membrane lipid phosphatidylethanolamine (PE) mitigated OPA toxicity through reducing cellular PE levels in cancer cells. In this study by Chidley *et al*, the authors showed that OPA forms pyrrole-adducts with the phosphoethanolamine headgroup leading to membrane destabilization and cancer cell death ^26^. While these studies all support the premise that OPA exhibits interesting anti-cancer activity that stems from alterations in mitochondrial function and lipid metabolism, the direct proteome-wide mapping of OPA protein targets remain understudied. OPA possesses multiple electrophilic moieties that can covalently react with nucleophilic amino acids on proteins, including cysteines and lysines, and multiple MOAs may contribute to its cytotoxic effects in a context-dependent manner. In this study, we profiled the anti-cancer activity of OPA across a panel of cancer cell lines and then used covalent chemoproteomic platforms to map the proteome-wide cysteine and lysine reactivity of OPA in a highly OPA-sensitive lung cancer cell line. We demonstrate how OPA covalently targets several proteins including two targets involved in the electron transport chain, leading to impaired mitochondrial metabolism and energetics and anti-cancer effects.

## Results

To identify human cancer cell lines that are particularly sensitive to OPA, we screened 283 tumor cell lines of diverse origin. We found that the lung squamous cell carcinoma line NCI-H1703 showed the lowest 50 % growth inhibitory (GI_50_) value across the screened cell lines with a GI_50_ of 0.17 µM **(Figure 1b; Table S1)**. We subsequently performed a more thorough dose-response of OPA on NCI-H1703 cell proliferation in serum-containing media and compared these data to the effects on cell viability with the C-6 epimer of OPA and the anhydro C-6 epimer of OPA, 6-*epi-*OPA and anhydro-6-*epi*-OPA, respectively **(Figure 1c-1d)**. These data demonstrated that OPA is significantly more sensitive in impairing NCI-H1703 cell proliferation with an EC_50_ of 0.54 µM compared to 6-*epi*-OPA and anhydro-6-*epi*-OPA with EC_50_ values of 3.7 and 4.0 µM, respectively **(Figure 1d)**. These data are consistent with the large majority of biological studies involving ophiobolins wherein OPA often shows greater cytotoxicity than its epimeric and anhydro counterparts ^7^. Additionally, administration of OPA at concentrations that impaired cell proliferation did not affect total cellular PE levels as determined by phospholipidomics **(Figure S1)**.

To identify the direct targets and potential mechanisms of this natural product, we next used isotopic tandem orthogonal proteolysis activity-based protein profiling (isoTOP-ABPP) chemoproteomic profiling approaches to map the cysteine and lysine reactivity of OPA in NCI-H1703 cells using previously established cysteine and lysine-reactive probes and methods ^31–34^. Across >2900 cysteines and >1000 lysines profiled, we identified 8 cysteine targets and 2 lysine targets that showed control versus OPA-treated probe-modified peptide ratios of greater than 2 with an adjusted p-value of <0.05 indicating significant engagement of these targets by >50 % **(Figure 1e-1f)**. These targets included C374 of farnesyl-diphosphate farnesyltransferase 1 (FDFT1), C100 of Ribosomal protein S6 (RPS6), C63 of heterogenous nuclear ribonucleoprotein A3 (HNRNPA3), C247 of geranylgeranyl diphosphate synthase 1 (GGPS1), C29 of Ras-related protein (RAB21), C511 of 2-hydroxyacyl-CoA lyase 1 (PHYH2), C127 of triosephosphate isomerase 1 (TPI1), C53 of mitochondrial HIG1 domain family member 2A (HIGD2A), K72 of cytochrome c oxidase subunit 5A (COX5A), and K292 of beta-actin-like protein 2 (ACTBL2) **(Figure 1e-1f, Table S2, and Table S3)**. These targets spanned several different protein classes and pathways, including FDFT1 in the cholesterol biosynthesis pathway, GGPS1 in the protein prenylation and geranylation pathway, TPI1 in glycolytic metabolism, and COX5A and HIGD2A in complex IV of the mitochondrial electron transport chain. Interestingly, COX5A and HIG2DA are both part of the same protein complex in the electron transport chain. We hypothesized that OPA may be disrupting mitochondrial metabolism and function through direct targeting of COX5A and HIG2DA.

We first sought to biochemically confirm the interactions of OPA and their derivatives with the identified targets using gel-based ABPP approaches competing OPA or OPA derivative binding against a rhodamine conjugated cysteine or lysine-reactive probe labeling of pure protein **(Figure 2a-2f, Figure S2)** ^32,35^. OPA showed dose-dependent binding to HIG2DA and COX5A via competition against cysteine and lysine-reactive probe labeling, respectively **(Figure 2a-2f)**. We also demonstrated OPA binds to RAB21, FDFT1, and GGPS1 **(Figure S2)**. Interestingly, 6-*epi-OPA* and anhydro-6-*epi-*OPA did not bind, or bound poorly, to HIGD2A and COX5A compared to OPA **(Figure 2a-2f)**. These results were consistent with our cell viablity data showing that 6-*epi-*OPA and anhydro-6-*epi-*OPA were less inhibitory to NCI-H1703 cell proliferation compared to OPA. We next used an alkyne-functionalized OPA probe (OPA-alkyne) to demonstrate direct, covalent, and dose-dependent labeling of pure COX5A and HIG2DA protein which was outcompeted by excess OPA **(Figure 2g-2i)**^28^. To further demonstrate target engagement in cells, we also showed enrichment of HIG2DA and COX5A from NCI-H1703 cells treated with the OPA-alkyne probe compared to vehicle-treated controls, upon subsequent copper-catalyzed azide-alkyne cycloaddition (CuAAC) with an azide-functioned biotin enrichment probe, avidin pulldown, and blotting for targets **(Figure 2j-2k)**. We further noted that unrelated proteins such as GAPDH were not pulled down with OPA-alkyne **(Figure 2j-2k)**.

**Figure 2.**
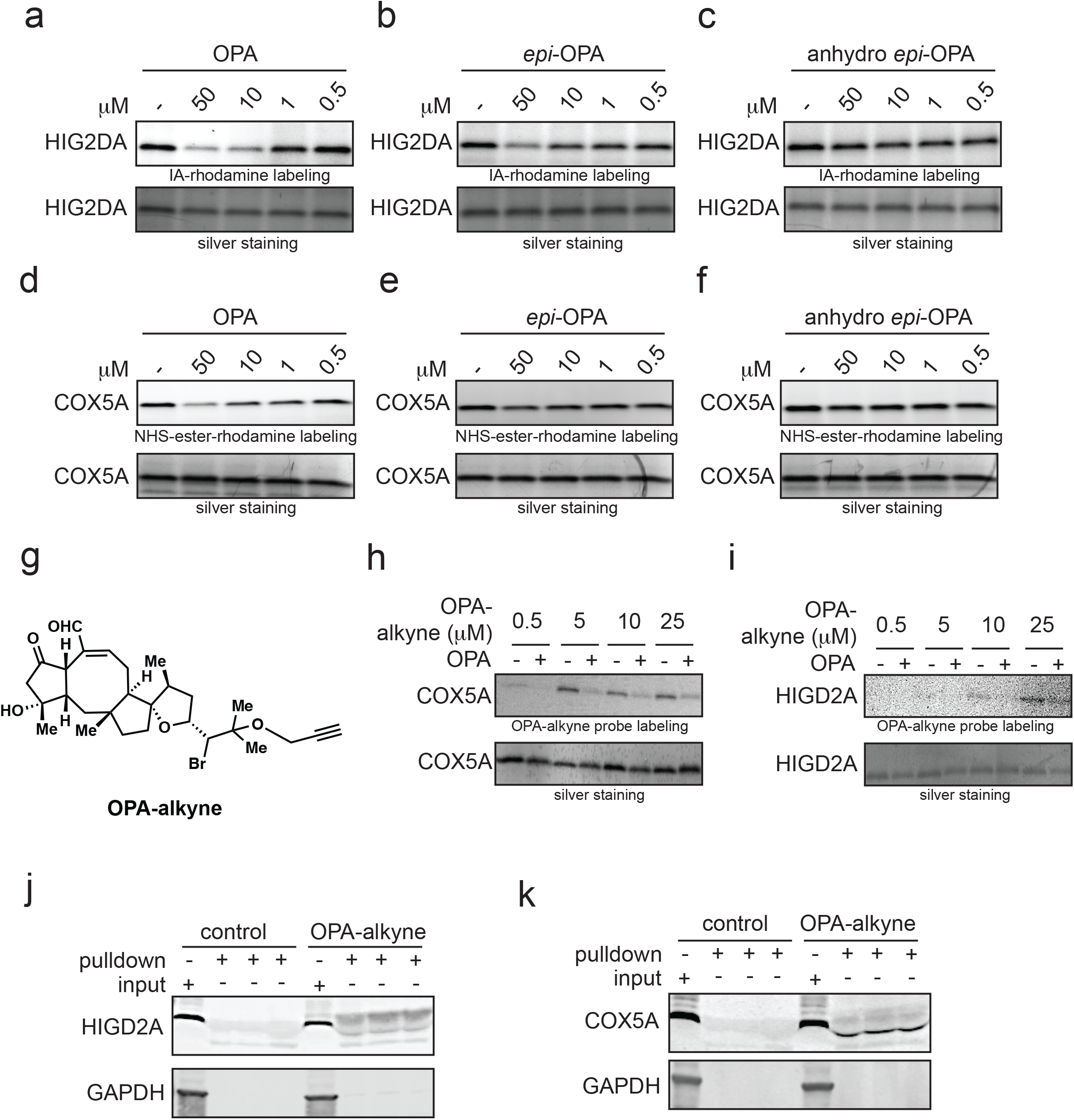
Biochemical characterization of OPA interactions with HIGD2A and COX5A. **(a-f)** Competitive gel-based ABPP analysis of OPA, 6-*epi*-OPA, and anhydro-6-*epi*-OPA binding to HIG2DA and COX5A. Pure proteins were pre-incubated with either DMSO, OPA, 6-*epi*-OPA, or anhydro-6-*epi*-OPA (30 min) prior to labeling of protein with either a cysteine-reactive rhodamine-functionalized iodoacetamide (IA-rhodamine) probe or lysine reactive rhodamine-functionalized NHS-ester probe (NHS-ester-rhodamine) probe. HIGD2A was labeled with 0.5 µM IA-rhodamine and COX5A labeled with 10 µM NHS-ester rhodamine. **(g)** Structure of alkyne-functionalized OPA probe (OPA-alkyne). **(h, i)** OPA-alkyne labeling of COX5A and HIG2DA. HIG2DA and COX5A pure protein (0.2 µg) were pre-incubated with DMSO vehicle or OPA (50 µM) for 30 min prior to labeling with OPA-alkyne probe for 60 min. **(j, k)** OPA engagement of HIG2DA and COX5A in NCI-H1703 cells. NCI-H1703 cells were treated with DMSO vehicle or OPA-alkyne (5 µM) for 1 h after which probe-modified proteins from cell lysate were appended to an azide-functionalized biotin enrichment handle for avidin enrichment and elution and blotted for HIGD2A or COX5A with GAPDH as the negative control GAPDH. Both input and pulldown protein levels are shown. Gels shown in **(a-f, h**,**i**,**j**,**k)** are representative of n=3 biologically independent replicates/group.

To understand the targets most responsible for the anti-proliferative effects of OPA, we next performed genetic validation studies. We demonstrated that individually knocking down the expression of COX5A, HIG2DA, RAB21, FDFT1, and GGPS1 all led to significant attenuation in OPA-mediated anti-proliferative effects **(Figure 3a-3d; Figure S2)**. Among these targets HIG2DA and COX5A knockdown both showed particularly significant resistance to OPA-mediated effects, indicating their larger importance among the targets **(Figure 3a-3d)**. Given that both HIG2DA and COX5A are in the same protein complex and knocking down one of these two targets may facilitate compensation by the other target, we also demonstrated significant attenuation of the anti-proliferative effects of OPA upon dual HIG2DA and COX5A knockdown **(Figure 3e-3g)**. Based on these genetic validation data, we placed the remainder of our focus on further investigating HIG2DA and COX5A as targets of OPA.

**Figure 3.**
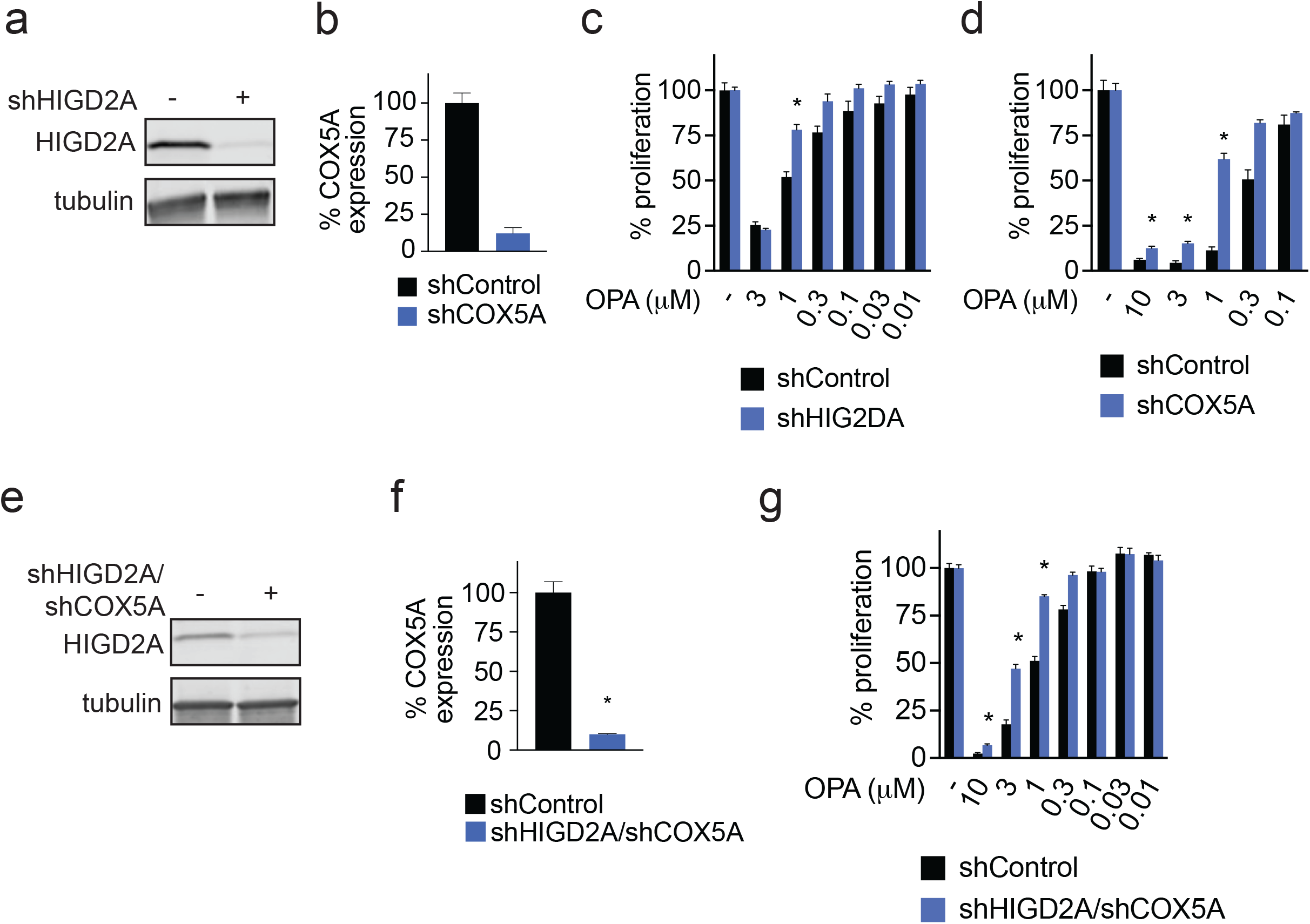
Genetic validation of OPA targets. **(a, b)** HIGD2A and COX5A stable short hairpin (shRNA) knockdowns in NCI-H1703 cells validated by Western blotting for HIGD2A **(a)** and by qPCR for COX5A **(b)** compared to shControl cells. **(c, d)** OPA effects upon cell proliferation in shControl, shHIGD2A, and shCOX5A NCI-H1703 cells. Cells were treated with DMSO vehicle or OPA for 24 h and cell proliferation was read out by Hoechst stain. **(e, f)** Validation of HIGD2A and COX5A dual and stable shRNA knockdown NCI-H1703 cells by Western blotting for HIGD2A **(e)** and qPCR for COX5A **(f)**. OPA effects upon cell proliferation in shControl versus dual shHIG2DA/COX5A NCI-H1703 cells. Cells were treated with DMSO vehicle or OPA for 24 h and cell proliferation was read out by Hoechst stain. Blots in **(a, e)** are representative of n=3 biologically independent replicates/group. Data shown are average ± sem in **(b, c, d, f, g)** and are n=3 **(b, f)** or n=6 **(c, d, g)** biologically independent replicates/group.

We next performed Seahorse metabolic studies monitoring mitochondrial oxygen consumption, a measure of mitochondrial electron transport chain activity. We showed that OPA activated mitochondrial oxygen consumption, suggesting that the targeting of HIG2DA and COX5A by OPA may activate these proteins in the complex IV of the electron transport chain **(Figure 4a)**. This enhancement in mitochondrial respiration was fully attenuated upon HIG2DA and COX5A knockdown and OPA in this case even has an inhibitory effect **(Figure 4b)**. This inhibitory effect may be through OPA interactions with another target that leads to inhibition of mitochondrial metabolism that is unmasked upon HIGD2A and COX5A knockdown. Based on these data, we hypothesized that OPA may over-activate mitochondrial electron transport and mitochondrial oxidative respiration leading to increased generation of reactive oxygen stress that would compromise the mitochondrial membrane potential and ATP generation, as has previously been shown with dysfunctional mitochondria and mitochondrial oxidative stress in cancer ^36^. Consistent with this premise, we observed an initial dramatic spike in mitochondrial ATP levels during the first hour of OPA treatment likely due to complex IV activation by OPA targeting of HIG2DA and COX5A **(Figure 4c)**. OPA treatment eventually led to significant impairment in mitochondrial membrane potential, more than known mitochondrial electron transport chain inhibitors or uncouplers **(Figure 4d)**. Subsequently, OPA treatment ultimately led to the collapse of mitochondrial oxidative phosphorylation, resulting in a near total loss of mitochondrial ATP production by 3 hours of treatment **(Figure 4c)**.

**Figure 4.**
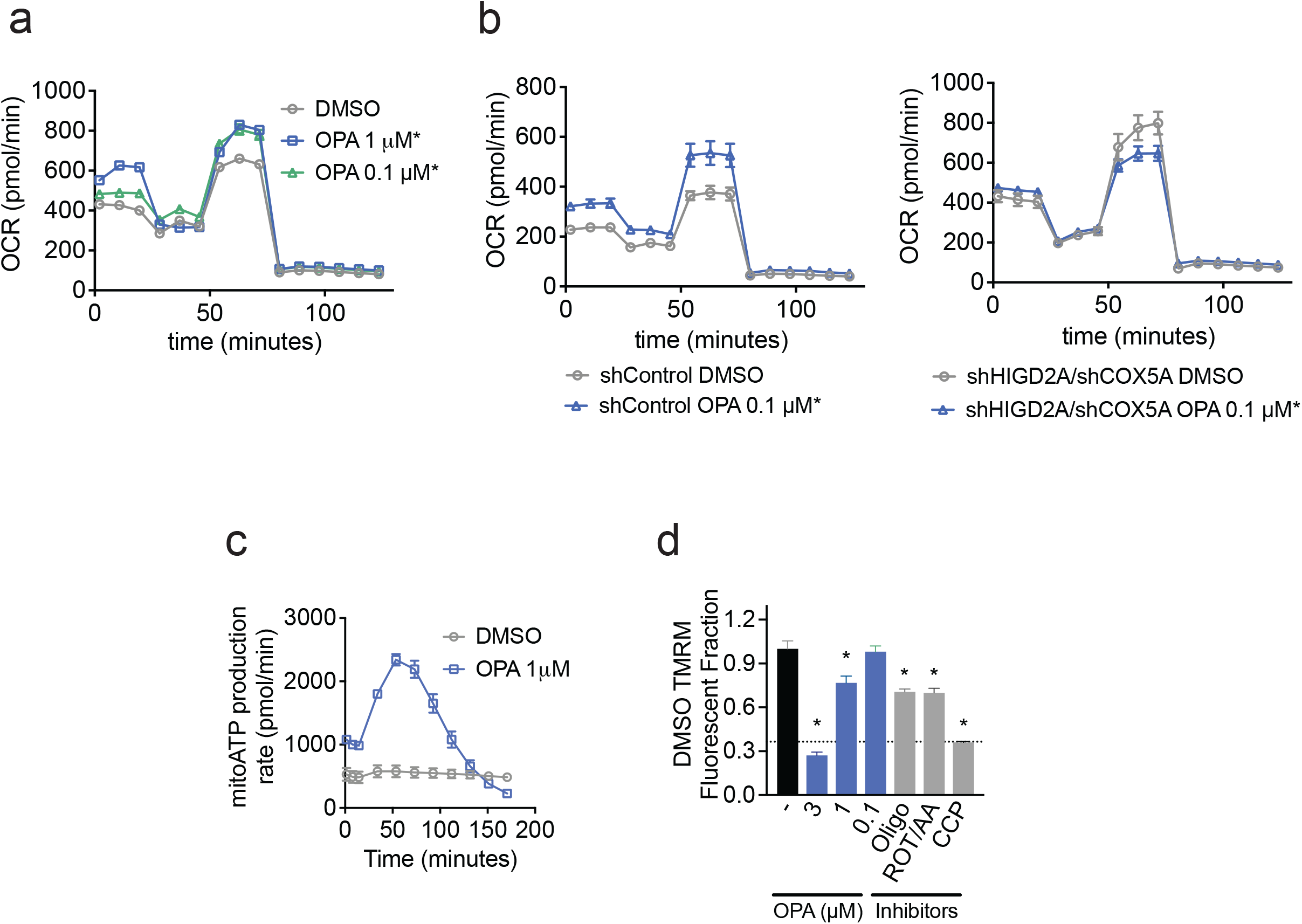
OPA effects upon mitochondrial oxygen consumption, ATP production, and mitochondrial membrane permeability. **(a, b)** Agilent Seahorse cell mitochondrial stress test results measuring oxygen consumption rate (OCR) in NCI-H1703 parental cells **(a)** and shControl versus shHIGD2A/COX5A dual knockdown **(b)** NCI-H1703 cells. Cells were pre-treated with either DMSO vehicle or OPA for 30 min prior measuring basal OCR for approximately 20 min prior to H^+^-ATPase inhibitor oligomycin (2.5 µM) treatment. ATP-linked respiration and proton leak were measured for approximately 20 min prior to mitochondrial uncoupler FCCP (2 µM) treatment. Maximal respiratory capacity recorded for 20 min prior to the experiment concluding with complex I inhibitor Rotenone and complex III inhibitor Antimycin A (0.5 µM) treatment. Data shown are average ± sem, n=7 biologically independent samples/group. **(c)** ATP production rate time-course study in NCI-H1703 cells treated with DMSO vehicle or OPA. Cells were treated with either DMSO vehicle or OPA 2.5 h prior to oligomycin (1.5 µM) treatment. Kinetic study concluded when Rotenone and Antimycin A (0.5 µM) were added. Data shown are average ± sem, n=7 biologically independent samples/group. **(d)** NCI-H1703 cells were treated for 2 h with OPA and other positive control mitochondrial inhibitors Oligomycin (2.5 µM), FCCP (2 µM), Rotenone/AA (0.5 µM), and CCP (10 µM), before being labeled with tetramethylrhodamine methyl ester (TMRM), a dye that accumulates in mitochondria that have active membrane potential, to measure mitochondrial membrane potential. Data shown in are in average ± sem, n=6 biologically independent samples/group.

## Discussion

In this study, we used covalent chemoproteomic platforms to map the proteome-wide cysteine- and lysine-reactivity of the electrophilic anti-cancer natural product OPA, revealing unique targeting of COX5A and HIG2DA within complex IV of the mitochondrial electron transport chain. This led to an over-activation of mitochondrial respiration, dissipation of mitochondrial membrane potential, and ultimately the collapse of mitochondrial metabolism and ATP production. Our genetic validation studies suggest that COX5A and HIG2DA are major contributors to OPA-mediated anti-proliferative effects and that the stimulation in mitochondrial metabolism is mediated through these targets as well. While there are many natural products, such as rotenone and oligomycin that inhibit the mitochondrial electron transport chain, our data point to OPA uniquely activating the mitochondrial electron transport chain leading to an unusual ATP spike, followed by a metabolic collapse. Our studies may also explain the mechanism underlying previous reports of OPA causing mitochondrial dysfunction ^20,29^. We note that our mechanism is likely one of several additional mechanisms that may underlie OPA effects, including previous reports that OPA inhibits human calmodulin ^18,30^, or that OPA can directly form pyrrole adducts with PE to destabilize membranes in certain cell lines ^26^. Taken more broadly, our study once again highlights the utility of using covalent ABPP chemoproteomic platforms with electrophilic and biologically active natural products to uncover unique mechanisms of anti-cancer activity ^37–39^, and open up the future possibility of exploiting such intriguing therapeutic modalities with more synthetically tractable chemical matter.

## Author Contributions

FAG, DKN, TJM conceived of the project idea, designed experiments, performed experiments, analyzed and interpreted the data, and wrote the paper. DQT, YW, BEP, DD performed experiments, analyzed and interpreted data, and provided intellectual contributions. JAT, JMK, MS provided intellectual contributions to the project and overall design of the project.

## Acknowledgement

We thank the members of the Nomura Research Group and Novartis Institutes for BioMedical Research for critical reading of the manuscript. This work was supported by Novartis Institutes for BioMedical Research and the Novartis-Berkeley Translational Chemical Biology Institute (NB-TCBI) for all listed authors. This work was also supported by the Nomura Research Group and the Mark Foundation for Cancer Research ASPIRE Award for DKN, FAG. This work was also supported by grants by the National Institutes of Health (R35CA263814 and R01GM116952). We also thank Drs. Hasan Celik, Alicia Lund, and UC Berkeley’s NMR facility in the College of Chemistry (CoC-NMR) for spectroscopic assistance. Instruments in the CoC-NMR are supported in part by NIH S10OD024998.

## Declaration of Interests

JAT, JMK, MS, and DD are employees of Novartis Institutes for BioMedical Research. This study was funded by the Novartis Institutes for BioMedical Research and the Novartis-Berkeley Translational Chemical Biology Institute. DKN is a co-founder, shareholder, and on the scientific advisory boards for Frontier Medicines and Vicinitas Therapeutics. DKN is a member of the board of directors for Vicinitas Therapeutics. DKN is also on the scientific advisory boards of The Mark Foundation for Cancer Research, Photys Therapeutics, Apertor Pharmaceuticals, Ecto Therapeutics, Oerth Bio, and Chordia Therapeutics. DKN is also on the investment advisory board of Droia Ventures and a16z Bio. TJM is a scientific advisory board member of Vicinitas Therapeutics.

## STAR Methods

### Chemicals

OPA was purchased from Millipore Sigma, Anhydro-6-*epi*-OPA was purchased from Cayman Chemicals. 6-*epi*-OPA was from previously published work ^12^. OPA alkyne probe was prepared according to known methods ^28^.

### Cell Culture

NCI-H1703 and HEK293T cells was obtained from UC Berkeley’s Biosciences Divisional Services Cell Culture Facility. NCI-H1703 cells were cultured in RPMI and HEK293T cells were cultures in DMEM media. Both media types contained 10%(v/v) fetal bovine serum (FBS), were supplemented with 1% glutamine, and were maintained at 37°C with 5% CO2. For shRNA knockdowns, cell specific media were supplemented with heat inactivated 10% FBS and with 1% glutamine.

### Cellular Phenotype Studies

Cell proliferation assays were conducted using Hoechst 33342 dye (ThermoFisher Scientific) as previously described. NCI-H1703 cells were seeded into 96-well plates (Corning 3904) at 25,000 cells per 150 µL of media and were left overnight to adhere. Cells were treated with 50µL of media containing 1:250 dilution from a 1000x DMSO compound stock and treated for 24hrs. After the incubation period, media with treatment was removed and 100 µL of diluted Hoechst 33342 dye in 10% formalin was added to each well and incubated at room temperature and in the dark for 15 minutes. Afterwards, the staining solution was removed and wells were washed with 100 µL of PBS twice. Prior to imaging, 100 µL of PBS was added to each well.

### In situ Competitive Mass Spec ABPP Chemoproteomic Studies

Competitive isoTOP-ABPP studies were done as previously reported ^34,40^. Cells were treated with either DMSO or 3µM of OPA for 4 hours. Each treatment group were harvested separately; cells were lysed by probe sonication in PBS and protein concentrations were measured by BCA assay. For both DMSO and OPA treated samples, 8 samples (at a concentration of 2 mg/mL) were aliquoted. Four DMSO and 4 OPA samples were treated with either 200 µM iodoacetamide-alkyne (IA-alkyne) probe or 500 µM lysine-reactive NHS-ester-alkyne probe and incubated for 90 minutes at room temperature. CuAAC was used by sequential addition of tris(2-carboxyethyl)phosphine (1 mM, Sigma), tris[(1-benzyl-1H-1,2,3-triazol-4-yl)methyl]amine (34 μM, Sigma), copper (II) sulfate (1 mM, Sigma), and biotin-linker-azide—the linker functionalized with a TEV protease recognition sequence as well as an isotopically light or heavy valine for treatment of control or treated proteome, respectively. After CuAAC, proteomes were precipitated by centrifugation at 6500 × g, washed in ice-cold methanol, combined in a 1:1 control/treated ratio, washed again, then denatured and resolubilized by heating in 1.2 % SDS/PBS to 80°C for 5 minutes. Insoluble components were precipitated by centrifugation at 6500 × g and soluble proteome was diluted in 5 ml 0.2% SDS/PBS. Labeled proteins were bound to avidin-agarose beads (170 μl resuspended beads/sample, Thermo Pierce) while rotating overnight at 4°C. Bead-linked proteins were enriched by washing three times each in PBS and water, then resuspended in 6 M urea/PBS (Sigma) and reduced in TCEP (1 mM, Sigma), alkylated with iodoacetamide (IA) (18 mM, Sigma), then washed and resuspended in 2 M urea and trypsinized overnight with 0.5 μg/μl sequencing grade trypsin (Promega). Tryptic peptides were eluted off. Beads were washed three times each in PBS and water, washed in TEV buffer solution (water, TEV buffer, 100 μM dithiothreitol) and resuspended in buffer with Ac-TEV protease and incubated overnight. Peptides were diluted in water and acidified with formic acid (1.2 M, Spectrum) and prepared for analysis. Isotopically light and heavy probe-modified peptides were analyzed by two dimensional LC-LC-MS/MS using a Thermo qExactive plus MS.

### Mass Spectrometry Analysis

Peptides from all chemoproteomic experiments were pressure-loaded onto a 250 μm inner diameter fused silica capillary tubing packed with 4 cm of Aqua C18 reverse-phase resin (Phenomenex # 04A-4299) which was previously equilibrated on an Agilent 600 series HPLC using gradient from 100% buffer A to 100% buffer B over 10 minutes, followed by a 5 minutes wash with 100% buffer B and a 5 minutes wash with 100% buffer A. The samples were then attached using a MicroTee PEEK 360 μm fitting (Thermo Fisher Scientific #p-888) to a 13 cm laser pulled column packed with 10 cm Aqua C18 reverse-phase resin and 3 cm of strong-cation exchange resin for isoTOP-ABPP studies. Samples were analyzed using an Q Exactive Plus mass spectrometer (Thermo Fisher Scientific) using a 5-step Multidimensional Protein Identification Technology (MudPIT) program, using 0 %, 25 %, 50 %, 80 %, and 100 % salt bumps of 500 mM aqueous ammonium acetate and using a gradient of 5–55 % buffer B in buffer A (buffer A: 95:5 water:acetonitrile, 0.1 % formic acid; buffer B 80:20 acetonitrile:water, 0.1 % formic acid). Data was collected in data-dependent acquisition mode with dynamic exclusion enabled (60 s). One full MS (MS1) scan (400–1800 m/z) was followed by 15 MS2 scans (ITMS) of the nth most abundant ions. Heated capillary temperature was set to 200°C and the nanospray voltage was set to 2.75 kV.

Data was extracted in the form of MS1 and MS2 files using Raw Extractor 1.9.9.2 (Scripps Research Institute) and searched against the Uniprot human database using ProLuCID search methodology in IP2 v.3 (Integrated Proteomics Applications, Inc). Cysteine residues were searched with a static modification for carboxyaminomethylation (+57.02146) and up to two differential modifications for methionine oxidation and either the light or heavy TEV tags. Probe-modified isotopically light or heavy differential modifications will be searched against cysteine modifications using m/z +464.26597 and +470.29977 for IA-alkyne; and against lysine modifications using m/z +494.26013 and +500.27394 for NHS-ester alkyne. Peptides were required to be fully tryptic peptides and to contain the TEV modification. ProLUCID data was filtered through DTASelect to achieve a peptide false-positive rate below 5%. Only those probe modified peptides that were evident across two out of three biological replicates were interpreted for their isotopic light to heavy ratios. For those probe-modified peptides that showed ratios >2, we only interpreted those targets that were present across all three biological replicates, were statistically significant, and showed good quality MS1 peak shapes across all biological replicates. Light versus heavy isotopic probe-modified peptide ratios were calculated by taking the mean of the ratios of each replicate paired light vs. heavy precursor abundance for all peptide spectral matches (PSM) associated with a peptide. The paired abundances were also used to calculate a paired sample t-test p-value in an effort to estimate constancy within paired abundances and significance in change between treatment and control. P-values were corrected using the Benjamini/Hochberg method.

### Bacteria Culture

In-house bacterial stocks and bacterial glycerol stocks purchased from Milipore Sigma (TRCN0000045961, TRCN0000234702, SHC016, SHC216, TRCN0000048127, TRCN0000045788, and TRCN0000304161) were cultured in LB overnight at 200rpm at 37C.

### Plasmid Isolation

E. coli containing desired plasmids were pelleted, lysed, and neutralized using commercially available QIAprep Spin Miniprep’s (catalogue no. 27104) user manual. Plasmid elutes’ concentration were determined using Nanodrop quantification.

### shRNA Knockdown Studies

Prior to HEK293T transfection, 3ug of each lentiviral plasmids—gene of interest(s), psPAX2 (carries GAG, REV, polgenes), and pMD2G (carries VSVG pseudotyping gene)— were diluted in 750µL of Gibco™ Opti-MEM™ I Reduced Serum Medium (catalogue no.31985-062). Lipofectamine 2000 (catalogue no.11668027) was also diluted in 750µL Opti-MEM. Both dilutions sat undisturbed at room temperature for 30 minutes prior to being mixed together and added to a 15cm plate of HEK293T cells. The following day, media was removed and replenished with fresh media. 72 hours after transfection (infection day), the viral soup was collected from HEK293T cells using 10mL luer lock syringes and filtered through a 0.45µM filter. An equal volume of target cell line media was added. Polybrene (catalogue no. TR-1003-G) in 1:1000 dilution was added. This lentiviral mixture was then added to the target cells. Following day, media was removed and replenished. 48 hours after infection, cells were selected on puromycin for 72 hours.

### Western Blotting

Antibodies to HIGD2a (Milipore Sigma, HPA042715-100UL), Rab21 (Milipore Sigma, R4530-25UL), FDFT1 (Milipore Sigma, HPA008874-100UL), GGPS (Milipore Sigma, HPA029472-100UL), GAPDH (Proteintech Group Inc., 60004–1-Ig), and α-Tubulin (Cell Signaling Technology, 3873S), were obtained from various commercial sources and dilutions were prepared per recommended manufacturers’ procedures. Proteins were resolved by SDS/ PAGE and transferred to nitrocellulose membranes using the BioRad system (1704159).

Blots were blocked with 5 % BSA in Tris-buffered saline containing Tween 20 (TBST) solution for 1 hour at room temperature, washed in TBST, and probed with primary antibody diluted in recommended diluent per manufacturer overnight at 4°C. Following washes with TBST, the blots were incubated in the dark with secondary antibodies purchased from Ly-Cor and used at 1:10,000 dilution in 5% BSA in TBST at room temperature. Blots were visualized using an Odyssey Li-Cor scanner after additional washes. If additional primary antibody incubations were required the membrane was stripped using ReBlot Plus Strong Antibody Stripping Solution (EMD Millipore, 2504), washed and blocked again before being reincubated with primary antibody.

### Gene Expression RTqPCR

Total RNA was extracted from cells using Trizol (Thermo Fisher Scientific) and aqueous solution was processed using QIAgen RNeasy Mini Kit (catalogue no. 74104). cDNA was synthesized using Qiagen Quantitect Reverse Transcription (205311) and gene expression was confirmed by qPCR using the manufacturer’s protocol Power SYBR Green Master Mix 2X concentration (4368577) on the CFX Connect Real-Time PCR Detection System (BioRad). Primer sequences for Fisher Maxima SYBR Green were derived from Primer Bank. Sequences of primers are as follow:

COX5a Forward: GGC TTA GGG GAC TGG TTG TC COX5a

Reverse: CCG TAA GAG GGC TTG GCT AC

PPIA (Cyclophilin) Forward: CCC ACC GTG TTC TTC GAC ATT

PPIA (Cyclophilin) Reverse: GGA CCC GTA TGC TTT AGG ATG A

### Gel Based ABPP

Gel-based ABPP methods were performed as previously described^34^. Recombinant pure human proteins were purchased from Origene: COX5A (TP306046), HIGD2A (TP301223), FDFT1 (TP301392), GGPS1 (TP302699), and RAB21 (TP310510). Pure proteins (0.2 μg) were pre-treated with DMSO vehicle or covalently-acting small molecules for 30 minutes at 37°C in an incubation volume of 25 μL PBS, and were subsequently treated with Tetramethylrhodamine-5-iodoacetamide dihydroiodide (Setareh) (IA-rhodamine) for 1 h at room temperature in the dark. Samples were then diluted with 10 μL of 4 × reducing Laemmli SDS sample loading buffer (Alfa Aesar) and heated at 90°C for 5 minutes. The samples were separated on precast 4–20% TGX gels (Bio-Rad Laboratories, Inc.). Prior to scanning by ChemiDoc MP (Bio-Rad Laboratories, Inc), gels were fixed in a solution of 10% acetic acid, 30% ethanol for 2 hrs. Inhibition of target labeling was assessed by densitometry using ImageJ.

### In Situ OPA-Alkyne Probe Labeling and Biotin-Azide Pulldown

OPA-alkyne probe was prepared according to known methods ^28^. Experiments were performed following an adaption of a previously described protocol. NCI-H1703 were treated with either vehicle (DMSO) or 5μM OPA alkyne probe for 4hours. Cells were harvested in PBS and lysed by sonication. Following instructions if for a sample of n=1 (this procedure was done in n=3 for pull down validation and TMT analysis). For preparation of Western blotting sample, 195 μL of lysate (4 mg/ml) was aliquoted per sample to which 25 μL of 10% SDS, 5 μL of 5 mM biotin picolylazide (900912 Sigma-Aldrich) and 25 μL of click reaction mix (3 parts TBTA 5 mM TBTA in butanol:DMSO (4:1, v/v), 1 part 50 mM Cu(II)SO_4_ solution, and 1 part 50 mM TCEP). Samples were incubated for 2 hours at 37°C with gentle agitation after which 1.2 mL ice cold acetone were added. After overnight precipitation at −20 °C, samples were spun in a prechilled centrifuge at 1250 x g for 10 minutes allowing for aspiration of excess acetone and subsequent reconstitution of protein pellet in 200 μL PBS containing 1% SDS by probe sonication. At this stage, total protein concentration was determined by BCA assay and samples were normalized to a total volume of 230 μL, with 30 μL reserved as input. 20 μL of prewashed 50% streptavidin agarose bead slurry was added to each sample and samples were incubated overnight at room temperature with gentle agitation. Supernatant was aspirated from each sample after spinning at 90 x g for 2 minutes at room temperature. Beads were transferred to spin columns and washed 3× with PBS. To elute, beads were boiled 5 minutes in 50 μL LDS sample buffer. To elute, beads were boiled 5 minutes in 50 μL SDS sample buffer. Eluents were collected by centrifugation and analyzed by immunoblotting.

### Agilent Seahorse Metabolic Testing

80,000 cells per well were seeded in Agilent cell culture microplate and incubated in 37°C at 5% CO_2_ overnight. Probe sensitization, assay buffer preparation, and program settings were followed as suggested by Seahorse XF Cell Mito Stress Test Kit’s (catalogue number 103015-100) user guide. After the cells adhered, cells were treated with compound for 30 min. Oligomycin, FCCP, and Rotenone/ Antimycin A was added at a final concentration of 2.5 µM, 2 µM, and 0.5 µM, respectively.

### Real-Time ATP Rate Assay

40,000 cells per well were seeded in Agilent cell culture microplate and incubated in 37°C at 5% CO2 overnight. Probe sensitization, assay buffer preparation, and “induced” program settings were followed as suggested by Seahorse XF Cell Mito Stress Test Kit’s user guide (catalogue no. 103592-100). Oligomycin and Rotenone/ Antimycin A was added at a final concentration of 1.5 µM and 0.5 µM, respectively.

### Mitochondrial Membrane Potential Assay

Mitochondrial membrane potential was assessed using Mitoprobe TMRM (Thermo Fisher M20036). NCI-H1703 cells were seeded into 96-well plates (Corning 3904) at 25,000 cells per 150 µL of media and were left overnight to adhere. Cells were treated with 50 µL of media containing 1:250 dilution from a 1000x DMSO compound stock and treated for 2 h. After the incubation period, media was spiked with TMRM to have a final concentration of 150 nM and incubated for 30 min. Afterwards, the media with TMRM was removed and wells were washed with 100 µL of PBS twice. Prior to imaging, 100 µL of PBS was added to each well.

### Lipidomics

NCI-H1703 cells were seeded at a concentration of 2 × 10^6^ cells in 10 cm dishes overnight. The following day, cells were treated with either vehicle control or OPA for 4 hours at 37°C with 5% CO_2_. Cells were then harvested, pelleted, and stored on dry ice. Cell pellets were extracted in 3 ml of 2:1:1 chloroform:methanol:water with the addition of 1 nmoles of internal standards—dodecylglycerol (positive ionization mode) and pentadecanoic acid (negative-ionization mode). Phases were separated by centrifugation and bottom organic phase was collected, dried under a stream of nitrogen and subsequently the dried extract was resuspend in 120 µL of chloroform. This sample was transferred to a glass mass spectrometry vial for subsequent lipidomic analysis by single-reaction monitoring based LC-MS/MS using a 6460 Agilent QQQ-LC-MS/MS using previously established methods ^41^.

