## Supporting Information for "Ophiobolin A Covalently Targets Complex IV Leading to Mitochondrial Metabolic Collapse in Cancer Cells"

**Supplemental Table Legends**

**Table S1. Cancer cell line panel screening of OPA**

**Table S2. isoTOP-ABPP data for OPA cysteine-reactivity.**

**Table S3. isoTOP-ABPP data for OPA lysine-reactivity**

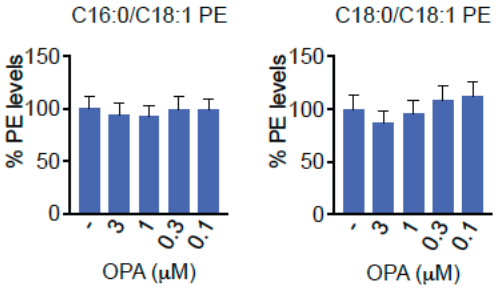


**Figure S1. Phosphatidylethanolamine levels in OPA-treated cells.** C16:0/C18:1 and C18:0/C18:1 phosphatidyl ethanolamine (PE) levels in NCI-H1703 cells treated with DMSO vehicle or OPA (3 μM) for 4h assessed by single-reaction monitoring (SRM)-based LC-MS/MS analysis. Data shown are average ± sem, n=4 biologically independent replicates/group. Related to **Figure S1.**

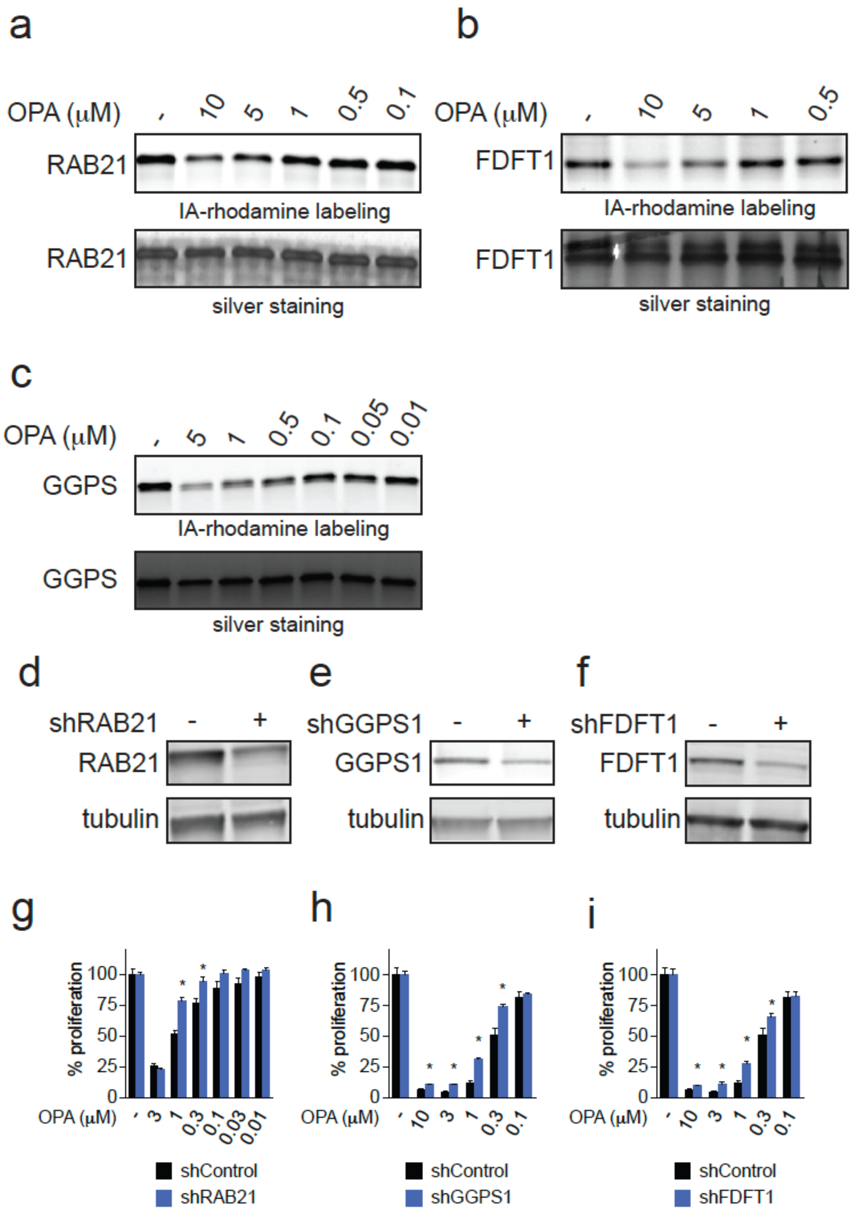


**Figure S2. Characterization of OPA targets. (a-c)** Competitive gel-based ABPP analysis of OPA binding to RAB21, FDFT1, and GGPS1. Pure proteins were pre-incubated with either DMSO or OPA (30 min) prior to labeling of protein with a cysteine-reactive rhodamine-functionalized iodoacetamide (IA-rhodamine) probe probe. RAB21 was labeled with 0.5 μM IA-rhodamine, FDFT1 labeled with 0.5 μM IA-rhodamine, GGPS labeled with 1 μM IA-rhodamine. Loading control is assessed by silver staining. Gels are representative of n=3 biologically independent replicates/group. **(d-f)** RAB21, GGPS1, and FDFT1 stable short hairpin (shRNA) knockdown in NCI-H1703 cells validated by Western blotting compared to shControl cells. **(g-i)** OPA effects upon cell proliferation in shControl, shRAB21 **(g)**, shGGPS1 **(h)**, or shFDFT1 **(i)** NCI-H1703 cells. Cells were treated with DMSO vehicle or OPA for 24 h and cell proliferation was read out by Hoechst stain. Blots in **(d-f)** are representative of n=3 biologically independent replicates/group. Data shown are average ± sem in **(g-i)** and are n=6 biologically independent replicates/group. Related to **Figure 2** and **Figure 3.**
